# Interrelationship of photoperiod and feed utilization on growth and reproductive performance in the Red eyed orange molly (*Poecilia sphenops*)

**DOI:** 10.1101/209346

**Authors:** Bela Zutshi, Aradhana Singh

**Affiliations:** Department of Zoology,Bangalore University, J.B. Campus, Bangalore. Karnataka, India, PIN. 560056; Research Scholar, Department of Zoology, Bangalore University, J.B. Campus, Bangalore. Karnataka, India, PIN. 560056

**Keywords:** Live-bearer Red eyed orange molly, Photoperiod, Growth parameters, Gonadal development, Fry production, Formulated feed

## Abstract

Light is a major environmental factors that synchronize all life-stages of fish, from body growth to sexual maturation. The objective of this research is to enhance the growth and reproductive performance in the test fish, red eyed orange molly (*Poecilia sphenops*) exposed to standardized manipulated photoperiod. In present study, growth and gonadal development of the test fish with an average initial weight (0.52g) and an average total length (3.2cm) exposed to two photoperiods (18L:6D and 10L:14D) at constant light intensity (1500 lx) for a period of 60 days was investigated. The fish were fed with formulated feed twice a day as per 10% of body weight. During a long day photoperiod (18L:6D) significant increase in growth parameters such as, weight gain (2.2 ± 0.04), feed conversion ratio (8.9 ± 0.004) and specific growth rate (1.2 ± 0% day^-1^) was observed. Mean gonadosomatic index (GSI) in female (30.4± 0.11) and male fish (2.0 ± 0.17) was also significantly higher in long-day photoperiod followed by short day photoperiod (10L:14D). The results confirm that manipulated long-day photoperiod induced somatic growth and enhanced gonadal development in *P. sphenops* without causing any stress.

**Summary:** This work is novel to research field since photoperiodic effect on *Poecilia sphenops*, a live-bearer fish under artificial environmental conditions resulted in enhanced growth and reproductive performance with fry production.

## 1. Introduction

Environmental factors induce metabolic changes in life activities of fish. Photoperiod is an important environmental factor that directly or indirectly influences fish growth, feeding, locomotor activity, metabolic rates, body pigmentation, maturation and reproduction (Gines *et al.*, 2004; Biswas *et al.*, 2005). Photoperiod and temperature are the best coordination for stimulating growth and reproduction by affecting internal timing system (Bromage *et al.*, 2001). El-Sayed and Kawanna (2004) studied the survival ability, weight gain, specific growth rate and feed conversion efficiency of fish Nile tilapia (*Oreochromis niloticus*) fry when exposed to 24L: 0D and 18L: 6D photoperiod regimes. The survival and growth of cultured fish depend upon the nutrients supplied and also the light available (Jeniffer *et al.*, 2012). According to Olivotto *et al.* (2003) feed conversion efficiency is affected not only through food intake but also by the exposure of fish to different light regime. Many fish species, including both marine and freshwater react to photoperiod regime and day length affecting growth. In recent years’ growth and gonadal development exposing Sea bass (*Dicentrarchus labrax*) for 40 days long photoperiod (12L:12D& 24L:0D), (Villamizar *et al.*, 2009), Nile tilapia (*Oreochromis niloticus L.*) for 24 weeks long photoperiod (24L:0D, 20L:4D & 18L:6D) (Rad *et al.*, 2006), Top mouth Gudgeon (*Pseudorasbora parva*) short and long photoperiod (0L:24D, 4L:20D, 8L:16D, 12L:12D, 16L:8D, 20L:4D & 24L:0D) for 90days (Zhu *et al.*, 2014). Several changes have been reported in growth, survival, reproductive patterns and levels of relevant hormones in fish species due to the effects of photoperiod regimes (Boeuf&Le Bail, 1999).

The *P. sphenops*, (*Poecilia* - many colored’ and ‘*sphenops*’ - ‘wedge appearance’) (red-eyed orange molly) a cyprinid fish belonging to Family - Poeciliidae is very demandable fish of the market share of ornamental fishes. This live - bearing, exotic and tropical ornamental fish commonly known as red eyed orange molly is found in stream, river and ponds of Mexico and United States. It is a tolerant species as it inhabits slight or moderate water streams, flood water ponds, lagoons, micro-reservoirs, lakes and reservoirs, and water ranging from clear to turbid, or even muddy (Miller *et al.*, 2005). The Molly comes in many different colors such as orange, black, green, gray and even white. The largest length registered for males and females was 96 mm and 83 mm, respectively. It is an omnivore but may eat commercial food such as pellets, algae, live or frozen food. It may survive in temperature 21°C - 28°C, pH: 7.5 - 8.5, Hardness: 10-25mg/l. The spawning season of *P. sphenops* occurs between July and October, concurring with the rainy season. Another reproduction peak registered is during February in natural conditions (Jose *et al.*,2016).

The gestation period for this fish is usually 28 days or more depending upon the photoperiodic regime. Fertilization is internal accomplished by insertion of gonopodium (highly modified anal fin) with milt in females (present only in males). Depending upon the size and maturity of fish which can be estimated by the length of the gonopodium, mollies produce broods of 25-40 live young, delivering them two to three times since females can store sperms long after mating. A single female gives birth multiple times throughout the year and has a lifespan of 5-6 years. Before the fry are born a dark triangle shaped patch around the anal vent known as ‘gravid spot’ showed the gravid condition of fish which becomes larger and darker as it matures as was also observed in the present study (Swain *et al.*, 2010). Sufficient data is not available regarding the ecology or general biology of molly under artificial environmental conditions.

Since ornamental fish farming has a high potential for profitability, the commercial production by the use of artificial photoperiods is widely accepted as tools for enhancing productivity of fish (Herve *et al.*, 2007). To achieve the complete benefit of reproductive performance and spawning frequency of fish in the aquarium and laboratory condition the environmental preferences of the fish needs to be understood. Environmental signals such as photoperiod and temperature are important inconsistent conditions for their growth and reproduction in laboratory condition. Thus, it is necessary to understand and evolve management techniques for their successful reproduction in captivity. A technique on growth and gonadal development studies has been cited on food fish and few ornamental fish. Studies on live-bearer fish, *P. sphenops* relating to the effect of photoperiod manipulation on their growth and gonadal development has not been reported so far since this finding will help the aqua-culturist to withstand the commercial demand. Thus the present study was conducted to fill this lacuna and to improve the reproductive management of the species in captivity of aquarium.

## 2. Materials and methods

### 2.1. Experimental fish

The experiment conducted with test fish *P. sphenops* (50no.) was obtained from its brood stock in Ornamental Fish Research Center, Hebbal, Bengaluru. The test fish with an average initial weight of 0.52g and average total length of 3.2cm was selected for the conduct of experiment.

### 2.2. Experimental design

To assess the objectives of the experiment, fishes were subjected to two artificial photoperiod regimes (18L:6D and 10L:14D) and natural light regime (control). The experiment was conducted in the laboratory using a fiberglass tank with a capacity of 50L for each experimental group. Water was aerated by a constant supply of air pump and 25% of water in each tank was renewed daily with fresh dechlorinated water so as to remove the feces and uneaten feed.

The physico-chemical characteristics of water were monitored every week. The dissolved oxygen (DO) ranged from 4.5 to 6.5mg/l, free ammonia also varied between 0.72 to 0.78mg/l (by standard method of APHA *et al.*, 2005) and pH (pH meter) values fluctuated as 7.0 to 7.5 in all the tanks throughout the experiment. Light was provided by a fluorescent lamp of 28W suspended about 40cm over the water surface in each photoperiod tank and light intensity was kept constant at 1500 lx in the experimental groups. The experimental tanks subjected to photoperiod regime were achieved by digital timer control.

### 2.3. Experimental procedures

Each experimental tank was stocked with 4 fish (1male: 3female) with three replicas for 60 days. Mean initial body weight (0.52-0.58g/fish) in the experimental groups and their replicates in each tank was stocked similar to avoid differences. The fish were acclimatized for 7days prior to start of the experiment and were fed with formulated feed used during the period of study. Prior to the experiment day fish were starved for 24 hours and total length and weight measured. Fish were fed twice per day (09:00h and 16:00h) with a formulated feed for the whole experimental period at the rate of 15% (28days), 10% (20days) and 5% (12days) of body weight per day for 60 days (Chapman, 2000). The detail of feed formulation in % of inclusion is as follows-

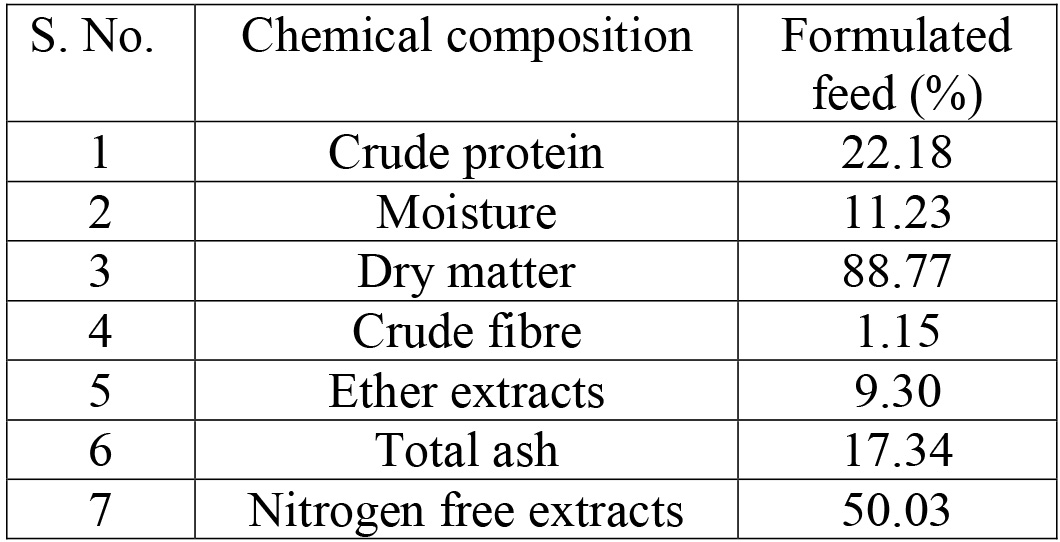
Composition of the formulated feed:

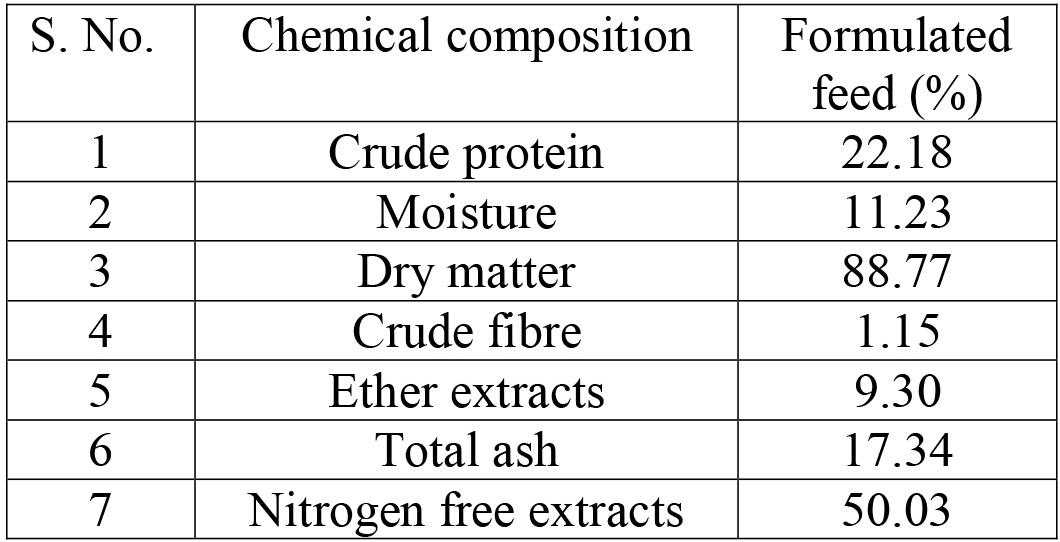
Proximate composition of formulated feed on dry matter basis (%):

### 2.4. Analysis for growth parameters

At the end of the experiment, the following growth parameters were analyzed:

Weight gain (WG), specific growth rate (SGR) and feed conversion ratio (FCR).
WG(%) = final body weight - initial body weight
SGR(%) = 100 × [L_n_ (W_2_)-L_n_ (W_1_)]/time (days)
Where, W_1_ and W_2_ indicate the initial and final weight (g), respectively.
FCR=Feed delivered to group/Live biomass gain of that group
Survival rate %=(Final fish number - Initial fish number) × 100/Initial fish number

### 2.5. Analysis of Gonadosomatic index (GSI)

At the end of the experiment fish were anesthetized by MS-222, and the gonads removed. For the female fishes the ovaries were weighed and opened to count the number of yolky eggs and embryos present recorded. For the male fish’s testes were weighed and gonadosomatic index (GSI) calculated by using the standard formula-

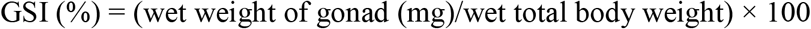

### 2.6. Data sampling and analysis

The data were analyzed and the results were expressed as mean (triplicates) ± SEM. The statistical significance of experimental and control groups was computed using one-way analysis of variance (ANOVA) with Tukey’s multiple comparison post-hoc test by using GraphPad Prism ver. 6.00 and the least significant difference was used to compare means at *P*< 0.05.

## 3. Results

All groups of fishes exposed to long (18L:6D) and short (10L:14D) photoperiod and also control ones were fed with formulated feed consisting of crude protein (22.18%), fish meal, ground oil cake, wheat flour, rice bran, minerals, vitamins & vegetable oil. The fishes were healthy and 0% mortality was observed during the experiment of 60 days. Growth performance and GSI of both male and female fish were analysed by standard methods. The number of fish with and without eggs and embryo were also recorded.

### 3.1. Growth performance

Growth patterns of *P. sphenops* exposed to artificial photoperiodic regimes in three replicates are presented in Fig. 1. Fish exposed to photoperiods, 18L:6D and 10L:14D for a period of 60 days revealed a significant increase in weight, feed conversion ratio and specific growth rate. The mean body weight gain showed significantly higher values (2.2 ± 0.04) in fish exposed to long photoperiod (18L:6D) (*P*<0.0001), when compared to those of short photoperiod (1.7 ± 0.06) and control (1 ± 0.06) (Fig.1A). Similar trend followed in mean FCR with its significant increase (8.9 ± 0.004) (*P*<0.0007) under exposure to 18L:6D, when compared to those of control (6.2 ± 0.004) (Fig.1B) as well as in mean SGR showing significant increase (1.2 ± 0.5% day^-1^) in fish exposed to 18L:6D and the lowest observed in control (0.8 ± 0.8% day^−1^) (Fig.1C). The difference between the mean SGR obtained under 18L:6D and 10L:14D photoperiod regime was not statistically significant (*P*>0.05).

**Fig. 1.**
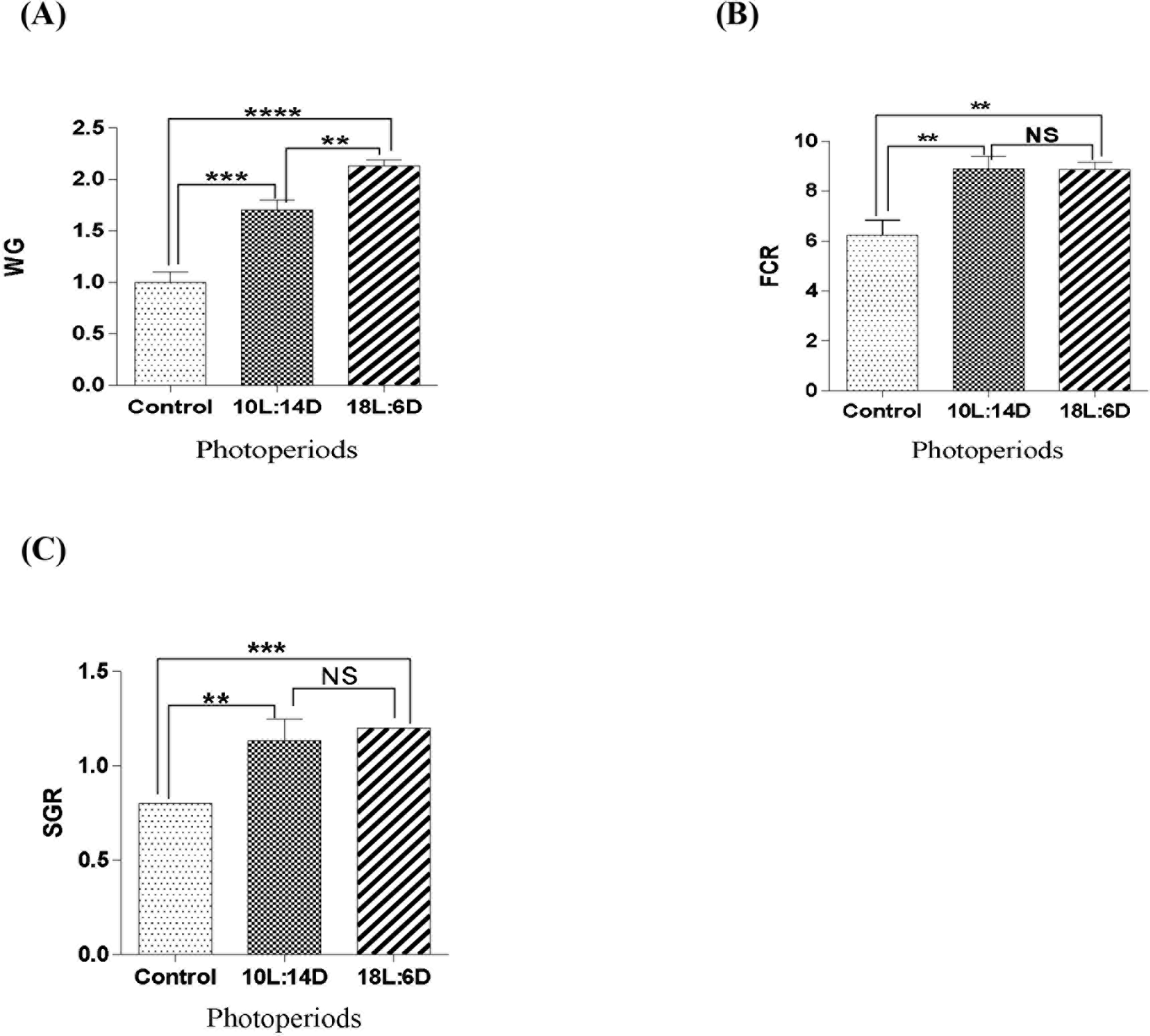
Effect of photoperiod on *P. sphenops.* Growth rate is represented in terms of (A) Weight gain (B) feed conversion ratio and (C) specific growth rate. Values are mean ± SEM of 4 fish/replica. Significance was calculated by one-way ANOVA and post-hoc test was done in accordance with Tukey’s comparison using GraphPad Prism 6 significance was represented as **** = *P*<0.0001; *** = *P*<0.001; ** = *P*<0.01 and NS= not significant

Growth performance (body weight) was noted every week for a period of 60days in the experimental and control groups. An increase in body weight after 28^th^day in the fish under long (18L:6D) and short (10L:14D)-photoperiod was observed and another significant rise in body weight exposed to (18L:6D and 10L: 14D) photoperiods after the 35^th^ day was noted (fig. 2). The fishes under control groups showed steady weight gain till the end of the experiment.

**Fig. 2.**
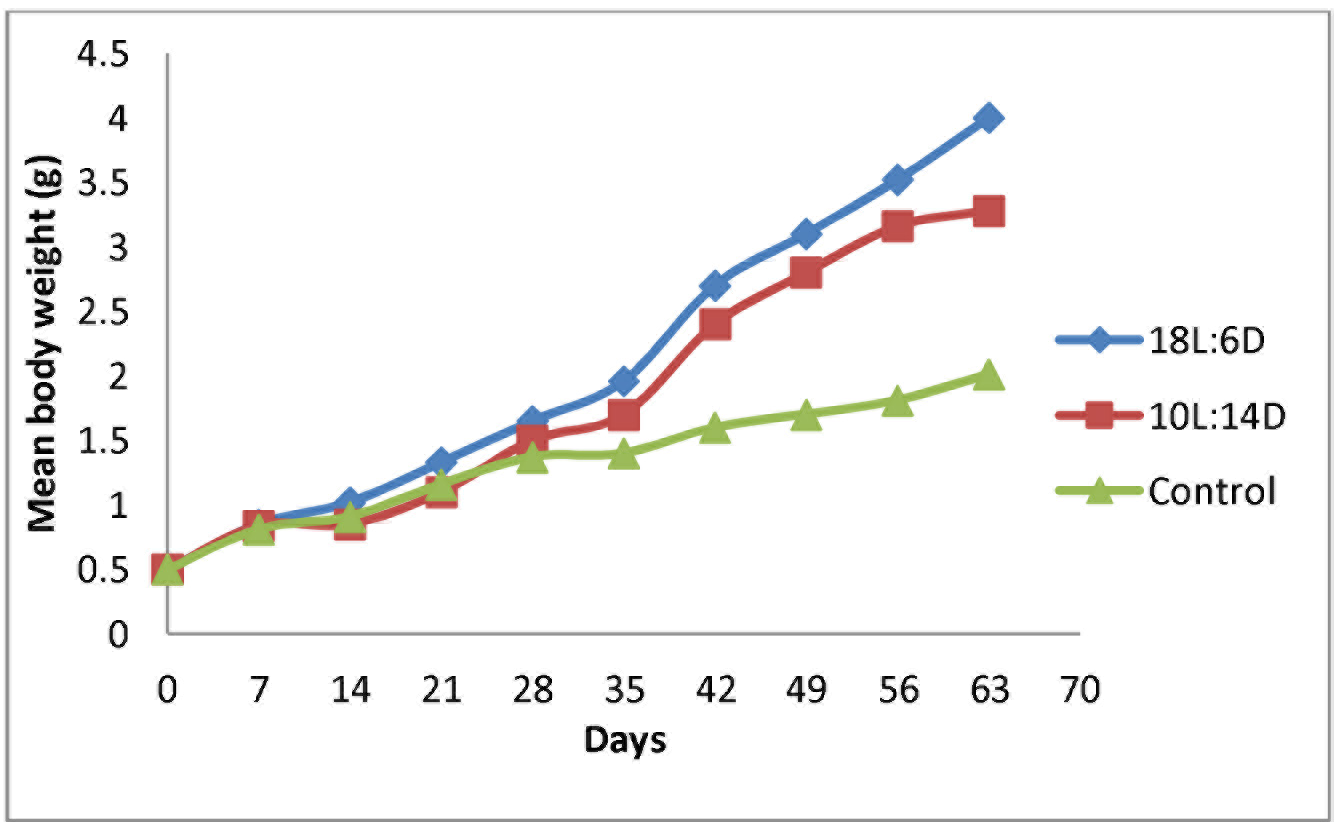
Weekly growth performance of *P. sphenops* under different photoperiods

### 3.2. Gonado-somatic indices (GSI)

Gonadal somatic index (GSI) of male and female *P. sphenops* showed significant changes during different photoperiod regime as represented in fig. 3. At the end of 60 days’ experiment significant differences were noted between mean GSI of female (*P*<0.03) and male fish (*P*<0.01) kept under a long photoperiod regime (18L:6D) and those in control group. A significant increase was recorded in mean GSI of male as well as female fish measuring 2 ± 0.17% and 30.4 ± 0.11% (respectively) exposed to 18L:6D photoperiod regime compared to those of control groups 1.0 ± 0.28 and 15.7 ± 0.03 (respectively). However, no significant difference was observed in mean GSI of male and female (1.9 ± 0.17% and 21.2 ± 0.03% respectively) exposed to a short photoperiod regime (10L:14D) compared to those under 18L:6D photoperiod (Fig. A & B). A Significant difference was observed within mean GSI of male and female fish maintained under 18L:6D and those of controls. Although the precise significance of the GSI of male fishes has not been established, it is likely that variation of this index reflects changes in spermatogenic activity.

**Fig. 3.**
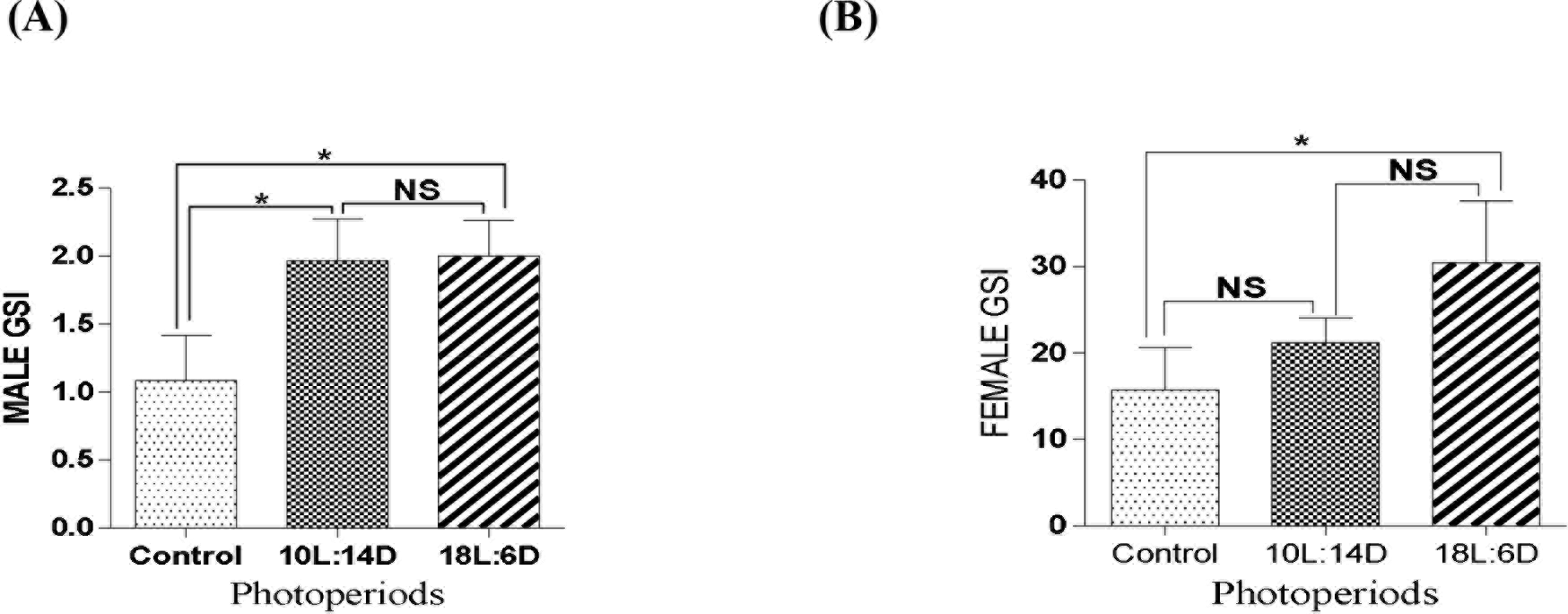
Effect of photoperiod on *P. sphenops*. Maturation is represented in terms of (A) male gonadosomatic index and (B) female gonadosomatic index. Values are mean ± SEM of 4 fish/replica. Significance was calculated by one-way ANOVA and post-hoc test was done in accordance with Tukey’s comparison using GraphPad Prism 6 significance was represented as *= *P*<0.02 and NS= not significant

**Fig. 1.**
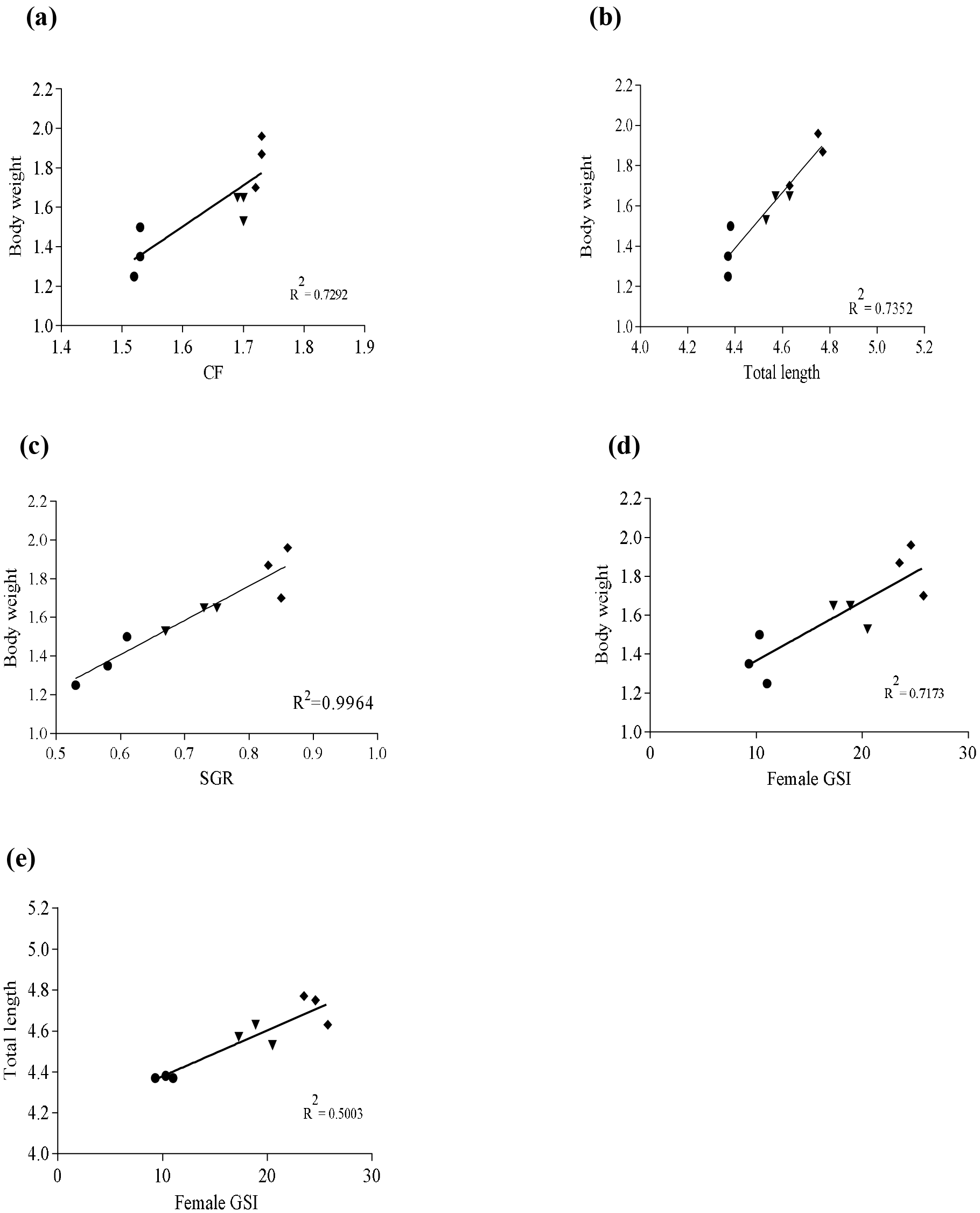
The linear relationship was assessed by linear regression and Pearson correlation coefficient using GraphPad Prism ver. 6.00, between body weight and (a) condition factor (CF) (b) total length (c) specific growth rate (SGR) (d) female gonadosomatic index (GSI), and (e) between total length and GSI of female fish. The curves were fitted by: (a) y = 2.084*X −1.832; (b) y =1.290*X −4.284; (c) y = 2.118*X + 0.1133; (d) y = 0.03029*X + 1.064 and (e) y = 0.01682*X + 4.267.

A significantly positive correlation was recorded between female fish body weight and gonad weight (*P*<0.01, *P*<0.04 & *P*<0.02). The regression relationship equations between body weight and gonads weight are shown in Table 1. A negative correlation between standard length and gonad weight was observed in all photoperiodic treated female fish and also in control groups (Table 2).

**Table 1.** Correlation between body weight and gonad weight of *P. sphenops*

| Treatment | Correlation | Regression | Significance value |
| --- | --- | --- | --- |
| Control | -0.6647* | $Y = -0.5009 * X + 1.101$ | $P<0.01$ |
| 10L:14D | 0.5949* | $Y = 0.4480 * X - 0.6997$ | $P<0.04$ |
| 18L:6D | 0.6576* | $Y = 1.081 * X - 2.176$ | $P<0.02$ |

Descriptive statistics for correlation and linear regression performed on body weight and gonad weight in *P. sphenops* and recorded positive correlation. There ‘*’ is representing significant value.

**Table 2.**
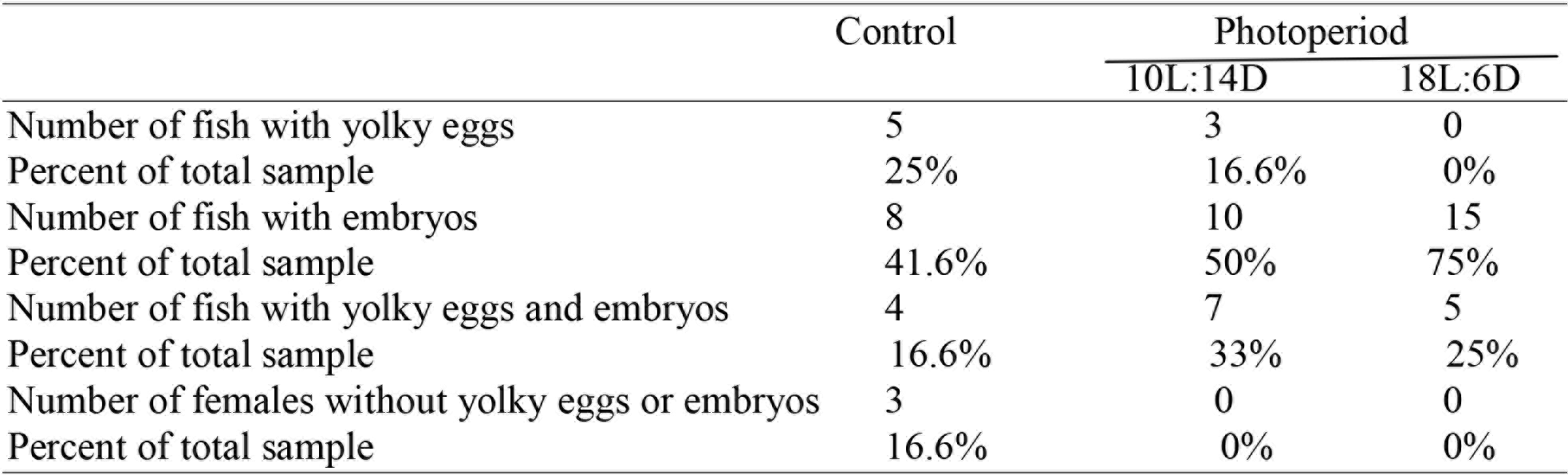
Correlation between standard length and gonad weight of *P. sphenops*

Descriptive statistics for correlation and linear regression performed on standard length and gonad weight in *P. sphenops* and recorded negative correlation. There ‘ns’ is representing no significant value.

### 3.3. Fishes with yolky eggs and embryos

Fishes exposed to long photoperiod showed significant number and percent of developed embryos, whereas those exposed to short photoperiod showed the maximum number of yolky eggs alongwith few embryos. Maximum numbers of fishes with yolky eggs (25%) were observed in control groups, whereas those exposed to long photoperiod revealed absence of yolky eggs. Fishes consisting of only embryos were more in number (75%) when exposed to long photoperiod whereas maximum number of fish with both yolky eggs and embryos together (33%) were observed when exposed to short photoperiod as when compared to control fish group. Female fishes without yolky eggs or embryos were observed in control groups only (16.6%) (Table 3). Male fish in all the experimental and control groups showed fully matured testis after 60 days of experiment fed with same formulated feed as those of female fish.

**Table 3.**
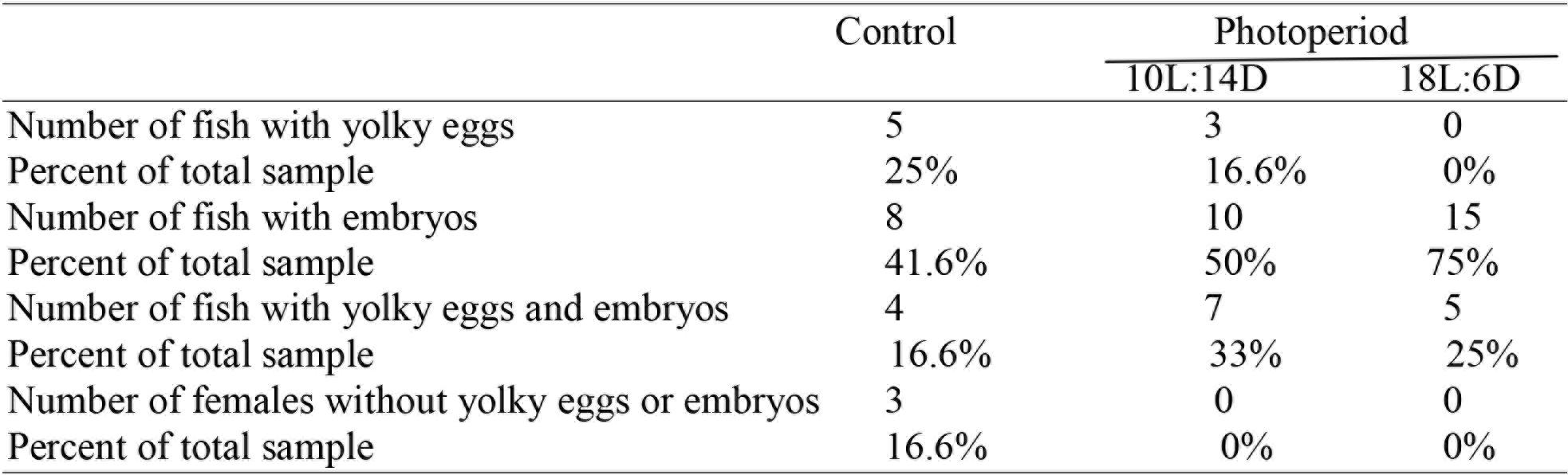
Effect of photoperiod on the reproductive performance of female *P. sphenops*

## Discussion

Photoperiod is an important physical factor which affects the growth, reproductive performance and maturation of the fish. Light and dark cycle provides internal harmonization for the rhythmic synthesis and release of hormones (i.e. melatonin), whose signal affects rhythmic physiological function in fish (Bairwa *et al.*, 2013). Optimization of environmental parameters and achieving the correct balance can result in better growth and early reproduction (Nancy *et al.*, 2013). In the present investigation male as well as female of *P. sphenops* exposed to 18L:6D and 10L:14D photoperiod regime for a period of 60 days showed a steady increase in the body weight till 28^th^ day. Significant growth observed on 35^th^ day and subsequent growth of *P. sphenops* for a period till the 60^th^ day exposed to two photoperiods (18L:6D and 10L:14D) can be corroborated to an improved growth rate of Nile tilapia exposed to long photoperiod (24L:0D; 20L:4D and 18L:6D) by Rad *et al.* (2006). Similar observations were reported by El-Sayed and Kawanna (2004), on improved weekly growth rate of Nile tilapia fry maintained under 24h and 18h long photoperiod.

This presumption has a lacuna for stimulation of growth and enhancement of reproductive performance in fingerlings of ornamental fish although it has been tested for species of food fishes. Since little is known about the exposure of male and female commercially important *P. sphenops* to manipulated photoperiod regimes with a specific time period to assess its effect on the growth and gonadal development along with the utilization of formulated feed, the present studies were conducted.

An increase in the final body weights, specific growth rates (SGR) and feed conversion ratio (FCR) of fish groups exposed to long day photoperiod (18L:6D) in the present investigation showed its positive correlation with hours of exposure to light since fish conserves more energy under longer photoperiod regime for growth patterns. The improved environmental rearing condition in laboratory including temperature, photoperiod and nutritional parameters like diet composition, ration and feeding frequency helps to gain best results in their culture for their growth and survival rate (Mohseni *et al.*, 2006). Responses of fish to photoperiodic manipulation stimulating growth with a simultaneous increase in FCR and SGR have been recorded by several authors, Rad *et al.* (2006) exposed Nile tilapia to artificial long-day photoperiods (24L:0D), 20L:4D and 18L:6D and Zhu *et al.* (2014) exposed *P. parva* to seven artificial constant-long day photoperiods resulting in a significantly higher weight gain in the fish exposed to 24L:0D<20L:4D<16L:8D than short-day (12L:12D<8L:16D<4L:20D) photoperiod regimes (p <0.05). Biswas *et al.* (2005) also reported that growth rate of *Oreochromis niloticus* L. can be enhanced significantly by using either long photoperiod (16L:8D) or continuous light regime (24L:0D). Shahkar *et al.* (2015), revealed similar observations in Caspian Roach showing an increase in weight and specific growth rate of fish exposed to continuous light photoperiod (24L:0D) compared to those exposed to 12L:12D, 6L:18D and 24D:0L. Fielder *et al.* (2002) reported increases in growth (total length) of the larva of snapper *Pagrus auratus*, exposed to long photoperiod 24L:0D and 18L:6D when compared to 12L:12D treatment. The significant increase in the growth rate of fish groups exposed to photoperiod (18L:6D) was accompanied by higher FCR and SGR can be due to the feeding strategy and type of feed reflecting closely to the maximum appetite. These findings are in agreement with the results of food fishes such as Nile tilapia larvae (ElSayed and Kawanna, 2004); Croaker miiuy (*Miichthys miiuy*) (Shan *et al.*, 2008); Nile tilapia fingerlings (Bezerra *et al.*, 2008; Cruz and Brown, 2009); Amazonian ornamental fish (*Pyrrhulina brevis*) (Veras *et al.*, 2013) and Persian sturgeon (*Acipenser persicus*) (Zolfaghari *et al.*, 2011). Improved appetite, greater food intake, higher feed conversion efficiency, higher digestibility and superior retention efficiency are the factors reportedly responsible for the faster growth rate under a long photoperiod regime (16L:8D). Increase in swimming activity of fish due to their exposure to more light hours 918L:6D) probably stimulated deposition of amino acids for the formation of muscle protein, leading to increased growth, since the deposition of protein is responsible for most of the weight gain when compared with other nutrients which constitute body composition (Biswas *et al.*, 2005).

Different species of fish at different stages of life respond to different photoperiods for their gonadal maturation, spawning and feeding rhythms (Mustapha *et al.*, 2014). Environmental factors such as light cues can have profound effects on the timing of gametogenesis, vitellogenesis and maturation in fish (Bromage *et al.*, 2001). Miranda *et al.* (2009) reported increases in GSI of Pejerrey fish, *Odontesthes bonariensis* when exposed to long photoperiod (18L:6D), regardless of the experimental temperature. Borg *et al.* (1987) suggested that in stickleback, *Gasterosteus aculeatus* species, long photoperiods induced maturation while shorter photoperiods did not do so even in high water temperature. This observation is in agreement with the present results in GSI values showing a gradual increase in gonadal weight and significant enhancement in body weight in fish exposed to long photoperiod (18L:6D) when compared to short photoperiod. GSI showed a minimum value in control fishes kept under laboratory conditions (control) than fish exposed to altered photoperiodic light regimes. The presence of fully matured, elongated and bulged testis in males and matured eggs and embryos in the ovary of female *P. sphenops* under a long photoperiod (18L:6D) parallels the findings in male and female Goldfish under the long light regime (19L:5D) (Sarkar and Upadhyay, 2011). Similar observations were reported by Zhu *et al.* (2014) in females of *P. parva* showing highest mean oocyte diameter exposed to long photoperiod (24L:0D) followed by 20L:4D and 16L:8D. Shirinabadi *et al.* (2013) also observed significantly high gonadal development in Whitespotted rabbitfish under 16 h long light regimes. GSI of the fishes exposed to an altered photoperiodic light regime showed higher values when compared to controlled ones. A positive correlation noted in the percentage increase in the gonad weight and body weight of male and female of *P. sphenops* exposed to photoperiod regimes makes it unlikely that variation in GSI value was due to fish weight alone, therefore the above mentioned result indicate that photoperiod is an important factor for regulating sexual maturation in *P. sphenops.* But the impact of longer photoperiod on growth and gonadal development still need to be explained by detailed studies related to hormonal aspect.

All the above mentioned data bring new information on the possibility to stimulate ornamental live-bearer fish growth and gonadal development by the different photoperiod regime. This finding will help the aqua-culturist to withstand the commercial demand since the fish *P. sphenops.is* one of the most preferred fish of aqua-culturist but poor availability in natural environment. Since it cannot be bred by them it becomes all the more expensive for the public and aqua-culturist. Discussion with the aqua-culturist in India requiring the Know-how of its breeding in tanks and maintenance at low cost feed led us to conduct the present study and to fill this lacuna and to improve the reproductive management of the species in captivity of aquarium was taken as a challenge. Thus experiments were conducted in laboratory conditions to enhance its growth performance, early maturation and captive breeding in tanks. Positive result was recorded for both sexes as mentioned in the paper.

An important finding that-the higher growth performance under manipulated photoperiods may be credited to the feed, feeding time as well as to improved activity and greater food intake’ is another factor that has led to the positive result. The manipulation of photoperiod did not cause any harmful and stressful response, and it is possible that gonadal development would be induced earlier than in the wild.

## Acknowledgement

This research did not receive any specific grant from funding agencies in the public, but Department of Zoology, Bangalore University Bengaluru is acknowledged for providing the necessary technical support.

